# *Wolbachia* infection in Argentinean populations of *Anastrepha fraterculus sp1*: preliminary evidence of sex ratio distortion by one of two strains

**DOI:** 10.1101/738500

**Authors:** CA Conte, DF Segura, FH Milla, AA Augustinos, JL Cladera, K Bourtzis, SL Lanzavecchia

## Abstract

**Background:** *Wolbachia,* one of the most abundant taxa of intracellular Alphaproteobacteria, is widespread among arthropods and filarial nematodes. The presence of these maternally inherited bacteria is associated with modifications of host fitness, including a variety of reproductive abnormalities, such as cytoplasmic incompatibility, thelytokous parthenogenesis, host feminization and male-killing. *Wolbachia* has attracted much interest for its role in biological, ecological and evolutionary processes as well as for its potential use in novel and environmentally-friendly strategies for the control of insect pests and disease vectors including a major agricultural pest, the South American fruit fly, *Anastrepha fraterculus* Wiedemann (Diptera: Tephritidae).

**Results:** We used *wsp*, *16S rRNA* and a multilocus sequence typing (MLST) scheme including *gat*B, *cox*A, *hcp*A, *fbp*A, and *fts*Z genes to detect and characterize the *Wolbachia* infection in laboratory strains and wild populations of *A. fraterculus* from Argentina*. Wolbachia* was found in all *A. fraterculus* individuals studied. Nucleotide sequences analysis of *wsp* gene allowed the identification of two *Wolbachia* nucleotide variants (named *w*AfraCast1_A and *w*AfraCast2_A). After the analysis of 76 individuals, a high prevalence of the *w*AfraCast2_A variant was found both, in laboratory (82%) and wild populations (95%). MLST analysis identified both *Wolbachia* genetic variants as sequence type 13. Phylogenetic analysis of concatenated MLST datasets clustered *w*AfraCast1/2_A in the supergroup A. Paired-crossing experiments among single infected laboratory strains showed a phenotype specifically associated to *w*AfraCast1_A that includes slight detrimental effects on larval survival, a female-biased sex ratio; suggesting the induction of male-killing phenomena, and a decreased proportion of females producing descendants that appears attributable to the lack of sperm in their spermathecae.

**Conclusions:** We detected and characterized at the molecular level two *wsp* gene sequence variants of *Wolbachia* both in laboratory and wild populations of *A. fraterculus sp.*1 from Argentina. Crossing experiments on singly-infected *A. fraterculus* strains showed evidence of a male killing-like mechanism potentially associated to the *w*AfraCast1_A - *A. fraterculus* interactions. Further mating experiments including antibiotic treatments and the analysis of early and late immature stages of descendants will contribute to our understanding of the phenotypes elicited by the *Wolbachia* variant *w*AfraCast1_A in *A. fraterculus sp.*1.

## BACKGROUND

*Wolbachia* constitutes a diverse group of maternally inherited endosymbionts belonging to the Alphaproteobacteria [1, 2]. To date, 16 different *Wolbachia* supergroups (A–F and H–Q) have been described [3]. Genomic approaches have been used to classify some of these *Wolbachia* supergroups as different species [4, 5], although this is still a rather controversial issue [6]. Supergroups A and B are widely spread across many arthropod taxa [7], C and D are found exclusively in filarial nematodes [8] whereas E is found in springtails [9]. Other *Wolbachia* supergroups are found in different host species. For instance, F supergroup comprises *Wolbachia* from termites, weevils, true bugs and scorpions [10, 11]. Different genetic markers have been employed to classify *Wolbachia* in supergroups including the 16S ribosomal RNA (*16S rRNA*) and the *Wolbachia* surface protein (*wsp*) genes [12–14]. More recently, two multi locus sequence typing (MLST) approaches and a *wsp*-based system have been developed for genotyping in addition to phylogenetic and evolutionary analyses of this bacterial taxonomic group [15, 16]. The discovery of supergroups (H-P) is mainly based on the full-length sequence of the *16S rRNA* and additional gene markers, such as *gro*EL (heat-shock protein 60), *glt*A (citrate synthase) and *fts*Z (cell division protein) [17]. This is in most cases associated to the lack of positive results (PCR amplification and/or sequencing) of any of the MLST genes in diverse supergroups (see also [3, 18–20]).

*Wolbachia* infections have been reported in the somatic tissues of a wide range of arthropod hosts [21, 22] and filarial nematodes [8, 23]. However, they are mainly known to reside in reproductive tissues and organs [21, 25–27]. In arthropods, *Wolbachia* often behave as reproductive parasites by manipulating the host reproduction to enhance its own vertical transmission [28] giving a reproductive advantage to infected individuals and spreading *Wolbachia* through natural populations [29–33]. A wide range of reproductive alterations induced by *Wolbachia* infection has been described in host species, including cytoplasmic incompatibility (CI), parthenogenesis, feminization and male-killing (MK) [1,2, 34–36]. CI is the most common phenotype induced by *Wolbachia* and is characterized by the induction of an embryonic lethality causing mating incompatibility in the crosses between *Wolbachia* infected males and uninfected females (unidirectional CI). Similar physiological incompatibilities are observed in crosses between individuals infected by mutually-incompatible *Wolbachia* strains (bidirectional CI) [35, 37–39]. Parthenogenesis is another well-documented *Wolbachia*-induced mechanism in haplo-diploid species by which the bacterium ‘forces’ unfertilized eggs to develop into females rather than males [40, 41]. *Wolbachia*-mediated feminization is characterized by the development of infected males into fertile females. This phenotype has been observed in both insects and isopods [42–44]. MK is expressed as male lethality during development resulting in a female-biased sex ratio [36, 45, 46]. MK can be elicited early during the embryonic development, or late in the larval or pupal stage [47]. MK is not limited to *Wolbachia*, as this phenomenon has been described for at least five clades of bacteria associated to the reproductive system (Additional File 1).

*Wolbachia*-host symbiotic associations are rather complex, since this reproductive microorganism can also be associated with a variety of additional phenotypes. These traits include the protection of insect hosts against pathogens and parasites [48–53], mating preference [54–56] and the response to olfactory cues [57]. The unique biology of *Wolbachia* has been explored for the development of novel strategies for the control of pests and diseases [33, 58–61]. For example, it has been shown that the Incompatible Insect Technique (IIT), which is based on the mechanism of *Wolbachia*-induced CI, can be used alone or in combination with the Sterile Insect Technique (SIT) to suppress populations of insect pests of agricultural, veterinary or human health importance [58, 62–67]. *Wolbachia*-induced MK has also been suggested as a tool for insect pest control [68, 69].

The South American fruit fly, *Anastrepha fraterculus* Wiedemann (Diptera: Tephritidae) is a complex of cryptic species [70–73] that is distributed in subtropical and temperate regions of the American continent, covering a wide geographical range from the United States of America to Argentina [74–76]. Recent studies focused on the elucidation of species from the *A. fraterculus* complex have followed an integrative approach. These scientific works addressed this taxonomic issue using different strategies based on morphology [73, 77], behavior and reproductive isolation [76, 78–81], and cytology and genetics [82–86]. Based on mating compatibility studies [87–89] and population genetic analysis [90, 91], a single biological entity of the *A. fraterculus* complex was identified in Argentina and southern Brazil. This taxon has been named *A. fraterculus sp.*1 by Selivon et al. [82] and Brazilian-1 morphotype by Hernández-Ortiz et al. [73]. The presence of *Wolbachia* has been described in Brazilian populations and in laboratory colonies of *A. fraterculus* from Argentina and Peru [79, 82, 92]. In addition, a recent publication [93] showed the presence of *Wolbachia* in *A. fraterculus* populations belonging to different morphotypes across America.

In the present study, we initiated a comprehensive study to detect and characterize *Wolbachia* infections in *A. fraterculus* from Argentina including a laboratory colony and three wild populations. After the detection and molecular characterization of the symbiont, we raised the hypothesis that *Wolbachia* infection may be associated with the induction of reproductive phenotypes, which could be a contributing factor in the speciation of *A. fraterculus* species complex. This hypothesis was tested with a series of crossing experiments assessing pre- or post-mating incompatibility, and these phenomena are discussed.

## MATERIALS AND METHODS

### Samples collection and DNA isolation

Wild *A. fraterculus* individuals were obtained from infested fruits collected in three different localities of Argentina: Horco Molle (Tucumán province); Villa Zorraquín (Entre Ríos province) and Puerto Yeruá (Entre Ríos province) (Table 1). Larvae and pupae obtained from each locality were maintained under standard laboratory conditions [94, 95] until emergence. In addition, individuals from the laboratory colony reared at IGEAF (INTA-Castelar, Buenos Aires, Argentina) were obtained, processed and stored under the same conditions until DNA extraction (Table 1). *A. fraterculus* IGEAF strain was established in 2007 with approximately 10000 pupae from the semi-mass rearing colony kept at Estación Experimental Agroindustrial Obispo Colombres, San Miguel de Tucumán, Tucumán, Argentina [96] and maintained to date (70 generations) under artificial rearing.

**Table 1.**
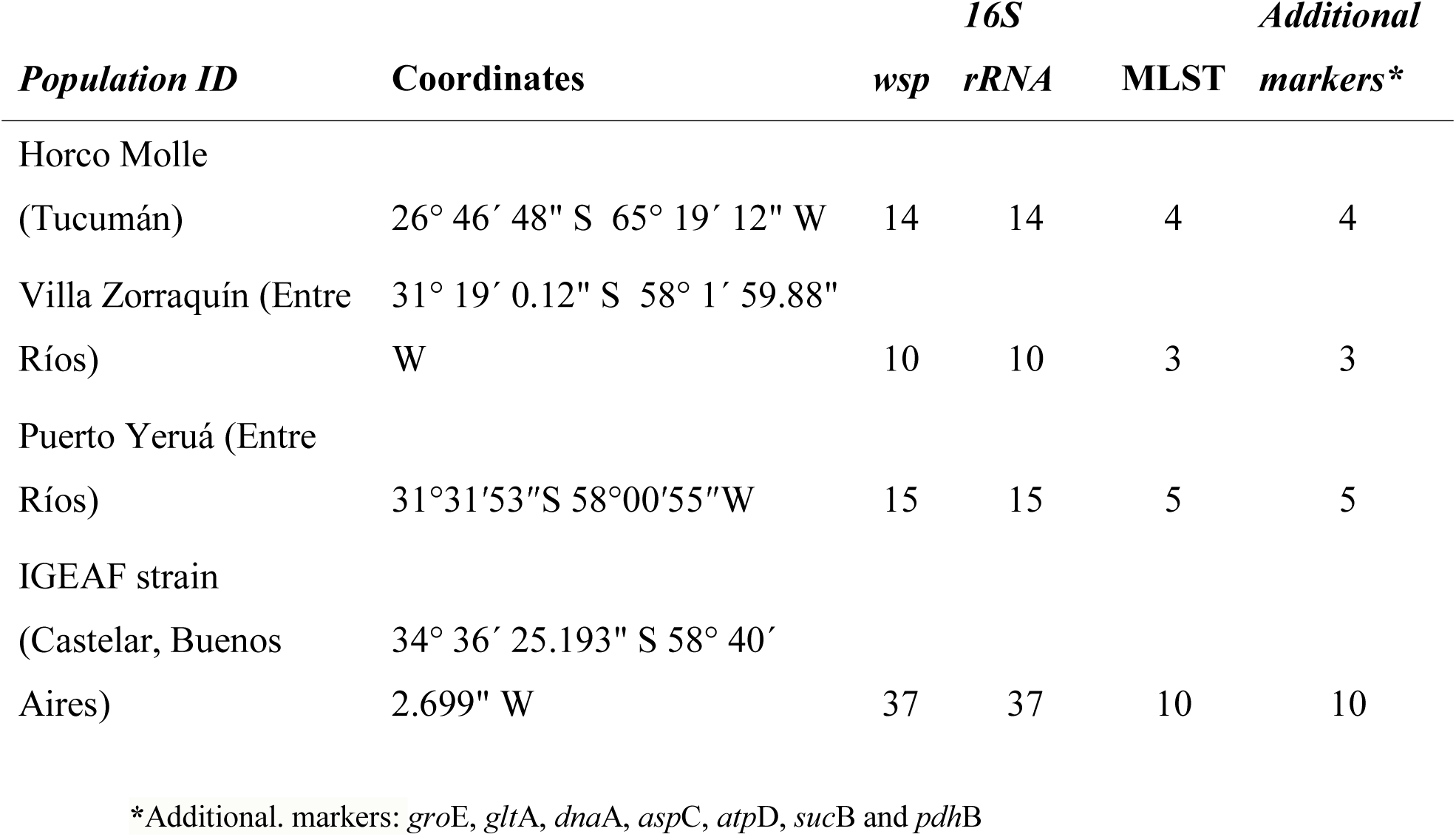
Sampling locations and number of individuals used for Wolbachia characterization.

All insects were washed with TE buffer (10 mM Tris–HCl, 10 mM EDTA, pH 8) and stored at −20°C until DNA extraction. Total DNA was individually isolated from adult flies (whole body) based on the protocol described by Baruffi et al. [97]. The quality of DNA samples was tested by electrophoresis in agarose gels 0.8 % w/v in buffer TBE 0.5 X and stained with ethidium bromide [98]. Images were captured with an UVP reveler (Fotodyne Inc. Hartland, WI, USA). Quality and quantity of DNA samples were also analyzed with Nanodrop 1000 (Thermo Scientific).

### Detection and genotyping of *Wolbachia* strains

*Wolbachia* detection was based on the amplification and sequencing of a *16S rRNA* gene fragment (438 bp) using the *Wolbachia*-specific primers wspecF and wspecR [99] and a *wsp* gene fragment (590 to 632 bp long) using primers 81F/691R [13]. The sequence characterization of a *wsp* gene from each *Wolbachia*-nucleotide variant found in this study was performed by *wsp* hypervariable regions (HVRs) analysis using the *Wolbachia* MLST database (pubmlst.org/*Wolbachia*). HVR alleles were determined based on comparisons among available translated nucleotide sequences [100]. Laboratory colony (37 individuals; 24 females, 13 males) and insects from natural populations (39 individuals; 22 females, 17 males) were analyzed. A subset of DNA samples (Table 1) were genotyped using the MLST scheme proposed by Baldo et al. [15] to characterize *Wolbachia*. Partial regions of *gat*B (aspartyl/glutamyl-tRNA(Gln) amidotransferase, subunit B), *cox*A (cytochrome c oxidase, subunit I), *hcp*A (conserved hypothetical protein), *fbp*A (fructose-bisphosphate aldolase) and *fts*Z genes were amplified, using the standard protocols provided in the *Wolbachia* MLST database [15]. PCR products were purified using a Wizard SV Gel and PCR Clean-Up System (Promega) and forward and reverse sequences were obtained using an Abi 3130XL Genetic Analyzer (Applied Biosystem, SIGYSA-INTA, Argentina). Sequences were manually edited and aligned using Bioedit 7.0.9.0 [101] and Staden Package [102].

A Neighbor-joining tree was reconstructed based on the concatenated MLST datasets (*gat*B, *cox*A, *hcp*A, *fbp*A and *fts*Z; 2,079 bases long) using sequences generated in the present study and a batch of representative nucleotide sequences belonging to A, B and D *Wolbachia* supergroups published by Baldo and Werren [103] available through the *Wolbachia* MLST webpage. The phylogenetic tree was constructed using Mega Version 5.1 software [104] based on the Jukes and Cantor [105] genetic distance model after 1000 bootstrap resamples.

Seven additional gene markers previously described for the genotyping of *Wolbachia* were utilized to distinguish *Wolbachia* genetic variants infecting the *A. fraterculus* Argentinean populations. Partial regions of *gro*EL and *glt*A [17], *dna*A (Chromosomal replication initiator protein) [106], *asp*C (aspartate aminotransferase) *atp*D (ATP synthase) *suc*B (dihydrolipoamide succinyltransferase) and *pdh*B (E1 component of the pyruvate dehydrogenase complex) [16] genes were amplified using primer sequences and PCR conditions described by the cited authors. At least three individuals of each *A. fraterculus* IGEAF strain harboring different genetic variants of *Wolbachia* were analyzed.

### Detection of other reproductive symbionts

*A. fraterculus* DNA samples were also screened for the presence of other reproductive symbionts (*Spiroplasma sp*., *Cardinium sp.*, *Rickettsia sp., Arsenophonus sp.* and *Hamiltonella sp*.) using the primers and conditions described by the authors cited in Table 2. In case of successful amplification, PCR products of expected size (according to the previously published works) were purified and sequenced.

**Table 2.**
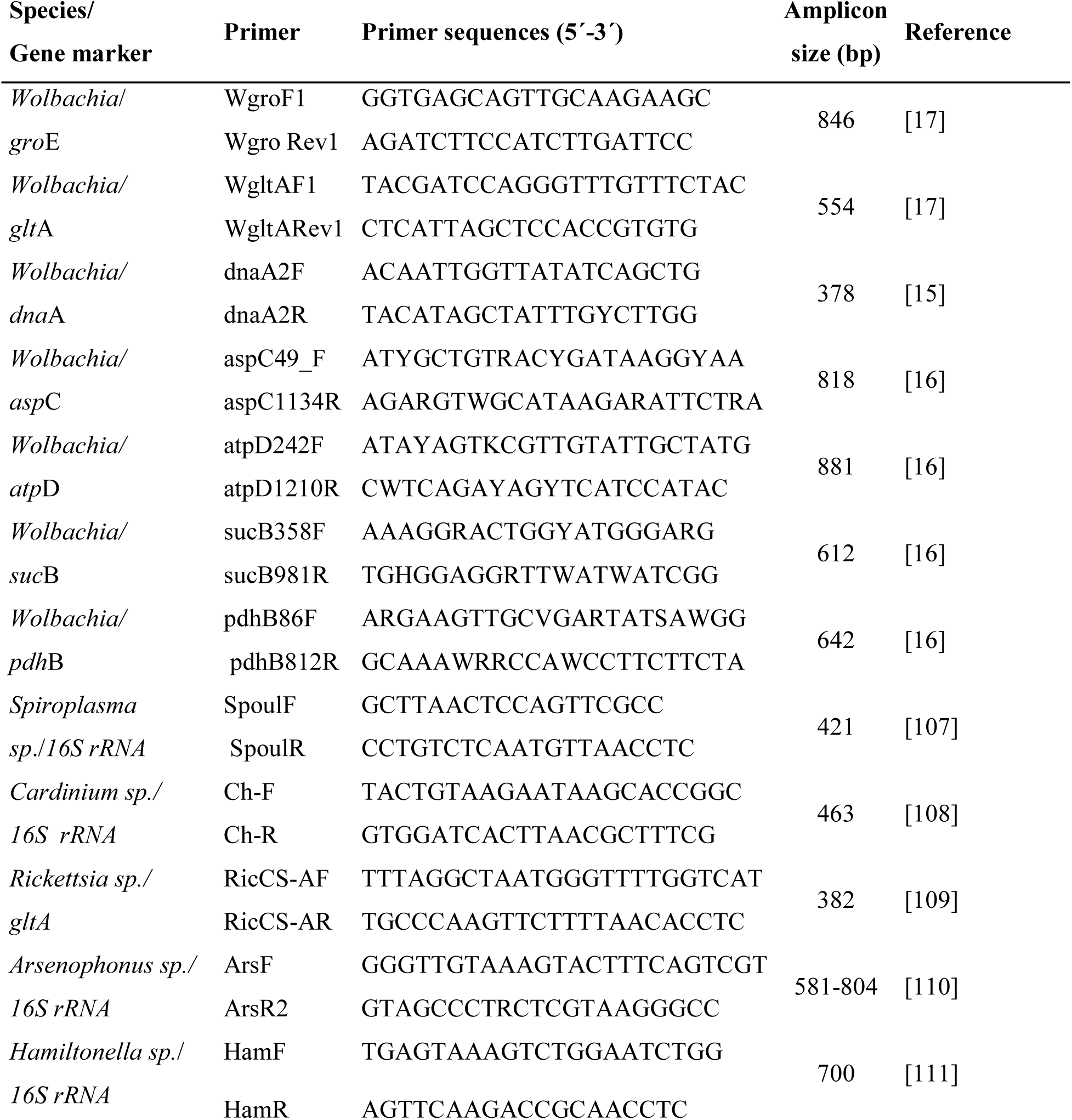
Additional primers used for the detection of *Wolbachia* and other symbionts.

New DNA sequences were deposited in public databases as is described in “Availability of data and material” section.

### Establishment of *A. fraterculus* Af-Cast-1 and Af-Cast-2 strains

At least 20 single pairs (female and male) from *A. fraterculus* IGEAF strain (IGEAF, INTA Castelar, Argentina) were maintained in standard conditions (25°C temperature; 50 % humidity and 12:12 Light: dark photoperiod) from the day of emergence to ensure that flies were virgin, since *A. fraterculus* reaches sexual maturity between 4 and 10 days after emergence [112]. At day 10 after emergence, egg collection devices (described by Vera et al. [94]) were offered to each couple continuously, either for a month or until at least 100 eggs were obtained. Total DNA was individually extracted from the parents of the families to determine the *Wolbachia wsp* nucleotide variant present in each one of them by PCR and sequencing of the amplicon as described above. Families sharing the same *Wolbachia* nucleotide variant (either *w*AfraCast1_A or *w*AfraCast2_A) were pooled and maintained as discrete strains under laboratory conditions of rearing. These *A. fraterculus* strains were named Af-Cast-1 and Af-Cast-2.

### Evaluation of *Wolbachia* genomic integration in *A. fraterculus*

The two laboratory strains of *A. fraterculus* (Af-Cast-1 and Af-Cast-2 strains) were treated with antibiotics. Eggs were deposited in plastic containers with larval diet [95] containing 0.01% rifampicin (Richet). After adult emergence, *Wolbachia* infection status was assessed by *wsp* and *16S rRNA* based PCR assays using the specific primers described above. DNA extracted from individuals of the Af-Cast-1 and Af-Cast-2 *A. fraterculus* strains reared without antibiotic treatment was used as a positive control.

Singly-infected *A. fraterculus* strains (Af-Cast-1 or Af-Cast-2) were maintained in our laboratory under standard rearing conditions [94].

### Mating experiments

In order to detect whether the presence of *Wolbachia* is associated with reproductive isolation, we carried out mating tests crossing *A. fraterculus* strains Af-Cast-1 and Af-Cast-2. Pre-zygotic isolation (which occurs before fertilization of gametes) as well as post-zygotic isolation (which occurs after fertilization) tests were performed as described below.

### Pre-zygotic isolation test

Individual crosses in every possible combination (i.e., female x male: Af-Cast-1 x Af-Cast-1, Af-Cast-1 x Cast-2, Af-Cast-2 x Af-Cast-1 and Af-Cast-2 x Af-Cast-2) were carried out in no-choice mating arenas under laboratory conditions following standard procedures [113]. Each arena consisted of a 1L plastic cylindrical container with a screen lid. The day before the test, 10 day-old (sexually mature) and virgin males were individually transferred to the mating arenas with no food or water. The next morning, under semidarkness, 15 day-old (sexually mature) and virgin females were released in the experimental arenas. Once the experiment was set up, the room lights were turned on (8:30 am). Experiments were conducted under laboratory conditions (T: 25 ± 1 °C and 70 ± 10 % RH). The number of replicates was 59±5 per cross type. The number of mated couples (percentage of mating), latency to mate and mating duration time were recorded for each type of cross. After the mating trial was completed, flies were removed from the mating arenas. Mated flies were preserved for post-zygotic tests (see below) whereas unmated flies were stored at −20 °C.

### Post-zygotic isolation test

Mated couples were maintained with food and water under controlled conditions and allowed to lay eggs in an artificial egg-laying device. Eggs were collected, placed on a piece of black filter paper, counted and transferred to Petri dishes (3 cm diameter) with larval diet [94, 95]. The Petri dishes were placed in a larger container on top of a layer of vermiculite (pupation substrate). After 5 days, the number of hatched eggs was recorded. After all developing larvae had exited the diet and pupated in the vermiculite pupae were collected, counted and placed under controlled conditions until emergence. The number and sex of emerged adults from each cross were recorded. Once the post-zygotic test ended, parental flies were stored at −20 °C and subsequently checked for the presence of *Wolbachia* (using the *wsp*-based PCR assay described above).

Ten F1 couples from each family (sibling mating) were randomly selected and kept under standard laboratory conditions with food and water and allowed to lay eggs to obtain F2, following the procedures described above for the parental generation.

### Data analysis

The percentage of mating recorded in the pre-zygotic test was compared among the four types of crosses by means of a chi-square test of homogeneity. The latency to mate and the mating duration time were compared among treatments using a one-way analysis of variance (ANOVA) followed by a post hoc Tukey’s multiple comparisons test.

Post-zygotic tests involved the analysis of the following parameters both in F1 and F2 generations: % of egg hatch (number of hatched eggs/total number of eggs*100); % of pupation (number of recovered pupae/number of eclosed larvae*100); % of adult emergence (number of emerged adults/number of recovered pupae*100); female sex ratio (number of adult females/number of emerged adults). These variables were analyzed by means of a one-way ANOVA. Normality and homoscedasticity assumptions were met for all variables, except for the percentage of pupation in the F1. In this case, data were arcsine square transformed to meet homogeneity of variances assumptions. In all cases, ANOVA were followed by post hoc Tukey’s multiple comparisons tests. Deviations from a 0.5 sex ratio were evaluated by means of a G-test of goodness of fit, applying the Bonferroni correction for multiple comparisons.

Additionally, we analyzed: 1. Percentage of mated females that produced eggs (number of females that laid > 10 eggs/number of mated females*100); 2. Percentage of females that produced viable eggs (number of females for which > 5% of eclosed eggs were found / number of females that produced eggs*100); 3. Percentage of females with descendants (number of females which produced > 5 emerged F1 adults/number of females that produced viable eggs *100); 4. Percentage of mated females that produced viable eggs (number of females for which > 5% of eclosed eggs were found/number of mated females*100; i.e., considering all mated females); 5. Percentage of mated females with descendants (number of females which produced > 5 emerged F1 adults/number of mated females *100; i.e., considering all mated females). These variables were compared among types of crosses by means of a Chi-Square test of homogeneity; first among the four types of crosses, and later between Af-Cast-1 and Af-Cast-2 females.

Statistical analyses were performed using STATISTICA for Windows [114].

### Cytological analysis

Mated females that did not produce descendants (females that did not lay eggs or that laid unviable eggs) were dissected under a stereoscope microscope (Olympus SZ30, Tokyo, Japan) to check both for any developmental abnormalities in the ovaries and the presence of sperm in spermathecae. The two ovaries and three spermathecae from each female were removed and placed on a slide. Preparations were stained with 2% acetic-orcein and observed under a phase contrast microscope Olympus BX40 (Olympus, Tokyo, Japan) using a 20X magnification objective. The general appearance, shape and structure of ovaries were analyzed as previously described [115, 116] and, the presence of sperm inside each one of the three spermathecae was visualized as previously described [112]. Sperm presence was determined whenever we visualized conspicuous bundles of sperm. For each female, the content of each spermatheca (presence/absence of sperm) was recorded.

## RESULTS

### Molecular characterization of *Wolbachia*

*Wolbachia* was positively detected in all the *A. fraterculus* adults tested (N=76; Table 1) using the *16S rRNA* and *wsp* gene PCR-based assays. *16S rRNA* sequence analysis showed identical base composition among samples (76 DNA samples, 380 bases). Basic Local Alignment Search Tool (BLAST) searches against the European Nucleotide Archive (ENA, EMBL, EBI) showed 100% identity with a large number of sequences including *w*Mel (*Wolbachia* endosymbiont of *Drosophila melanogaster*; GenBank accession DQ412083.1).

In the case of *wsp* gene sequences (507 bases) a unique non-synonymous nucleotide change (C/T) was detected among the 76 samples analyzed (Fig. 1). The *wsp* nucleotide variants detected were named *w*AfraCast1_A and *w*AfraCast2_A respectively. BLAST nucleotide search of *wsp* gene sequence from *w*AfraCast1_A showed 100 % identity with *A. fraterculus* isolate *w*AfBrazil_A (EU651897.1) and *A. fraterculus* isolate *w*AfPeru_A (EU651893.1) among others. The *wsp* nucleotide sequence of *w*AfraCast2_A showed 100% identity only with *A. fraterculus* isolate *w*AfArgentina_A (EU651896.1).

**Figure 1.**
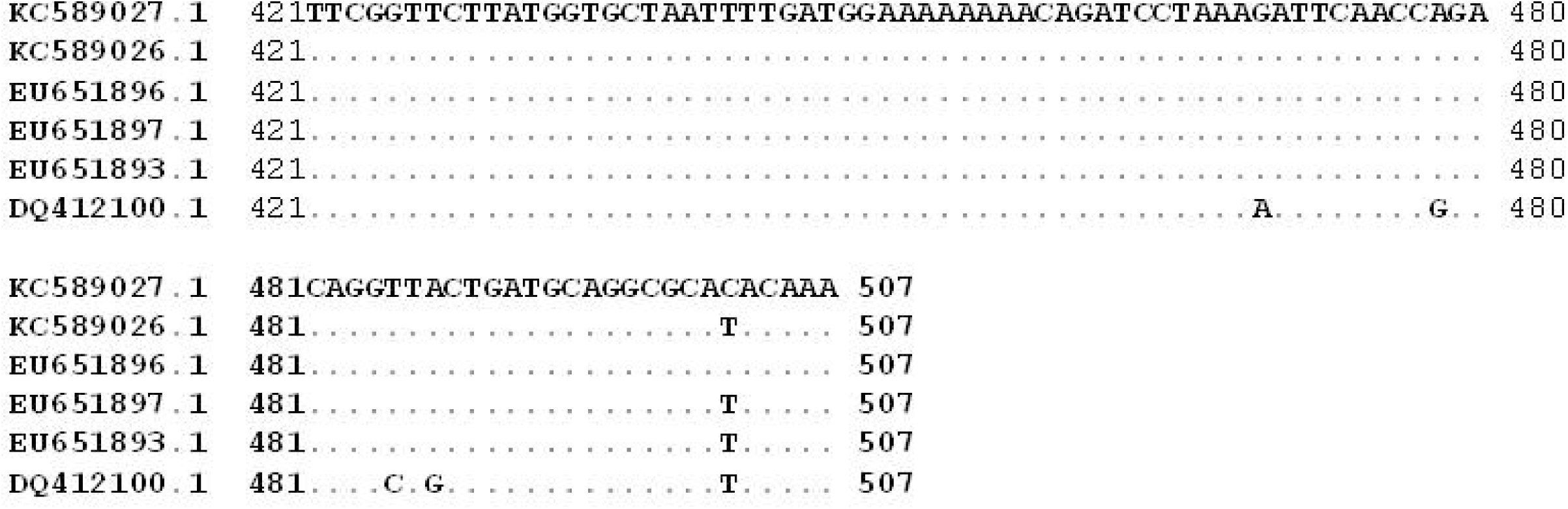
Identification of the single nucleotide substitution in 507 bp alignment of wolbachia wsp sequences. The figure shows a section of the *wsp* nucleotide sequences alignment including *Wolbachia* sequences described here (AN KC589026.1 and KC589027.1 GenBank) corresponding to wAfraCastl_A or wAfraCast2_A respectively and, sequences from GenBank (NCBI) corresponding to the *A.fraterculus* isolate wAfArgentina_A(EU651896.l);A. *fraterculus* isolate wAfBrazil_A (EU651897.l); *A.fraterculus* isolate wAfPeru_A (EU651893.1) and *Wolbachia* strain wMelinfecting *D. melanogaster* (DQ412100.1).

The analysis of the HVRs of the *wsp* gene performed through the *Wolbachia* MLST webpage, evidenced different *wsp* allele and allelic profiles in HVR4 for the *Wolbachia* nucleotide variants identified here (Table 3). Further HVRs allelic profiles comparison revealed perfect match between *w*AfraCast1_A and several *Wolbachia* strains including *Wolbachia* strains infecting *Rhagoletis cerasi* (Diptera: Tephritidae) and *Leucophenga maculosa* (Diptera: Drosophilidae), whereas *w*AfraCast2_A showed no perfect match in this database.

**Table 3.** Characterization of the *wsp* HVRs. HVR allele definition is based on the amino acid motifs analysis of the *wsp* gene sequence (61-573 bp) in respect of *w*Mel (*Wolbachia* databases - webpage pubmlst.org/*Wolbachia*/). Assigned alleles to *wsp* nucleotide sequences are also showed (*wsp* allele). of cross, and female proportion obtained in the offspring (F1 and F2).

| *Wolbachia* strain ID | HVR1 | HVR2 | HVR3 | HVR4 | *wsp* allele |
| --- | --- | --- | --- | --- | --- |
| *w*AfraCast1_A | 1 | 12 | 21 | 19 | 23 |
| *w*AfraCast2_A | 1 | 12 | 21 | 283 | 663 |

MLST analysis showed identical nucleotide sequences in 22 DNA samples from the different *A. fraterculus* populations evaluated (Table 1). The MLST allelic profile obtained corresponds to *gatB*:1, *coxA*:1, *hcpA*:1, *ftsZ*:3 and *fbpA*:1 and sequence type (ST) 13. Phylogenetic analysis based on a concatenated dataset of 5 MLST loci (2,079 bases) including the nucleotide sequences obtained here and a dataset of representative sequences from A, B and D *Wolbachia* supergroups from Baldo and Werren [103] revealed that *Wolbachia* found in Argentinean *A. fraterculus* populations belong to supergroup A (Fig. 2).

**Figure 2.**
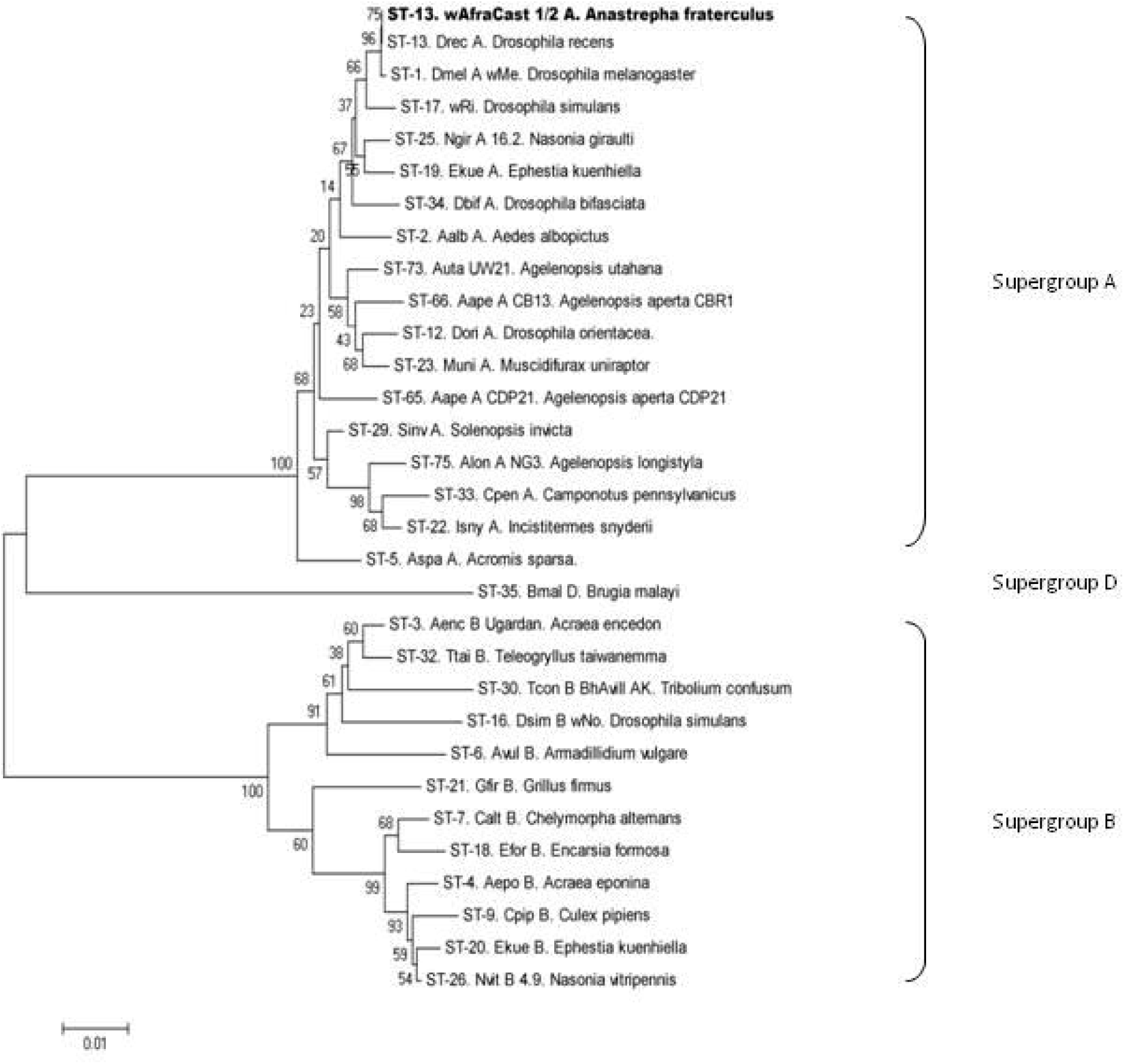
Neighbor-joining tree reconstructed based on concatenated MLST data (2079 bases). Phylogenetic tree reconstructed using a dataset including 30 MLST concatenated sequences published by Baldo and Werren [103] and a unique sequence corresponding to the concatenated MLST from wAfraCast l/2_A. Branch name is identified as *Wolbachia* sequence type (ST)-*Wolbachia* strain (if known)-host species name. Numbers in nodes indicate bootstrap support percentage (1000 replicates). *Wolbachia* supergroups are shown to the right of the tree. Similar topology was observed using Maximum Likelihood analysis (Additional File 4).

In addition to MLST analysis, we evaluated the polymorphisms in seven additional loci from the *Wolbachia* genome (*gro*EL*, glt*A*, dna*A*, suc*B*, asp*C*, atp*D and *pdh*B) in at least three individuals of Af-Cast-1 and Af-Cast-2 strains. After the analysis of at least 370 b from each locus (see details in Table 2) no polymorphism was identified showing a high similarity between *w*AfraCast1_A and *w*AfraCast2_A at genomic level (see sequence alignments in Additional File 2). The sequence comparisons using BLAST also evidenced similarities among sequences from *Wolbachia* infecting *Drosophila* species (*w*Mel, *w*Ri, *w*Ha) for the five genes evaluated, confirming the results obtained by MLST and phylogenetic analyses of *w*AfraCast1/2_A clustered with *w*Mel group from supergroup A (Fig. 2).

### Prevalence of *Wolbachia*

*Wolbachia* was detected in 100% of *A. fraterculus* individuals through PCR amplification and sequencing of *wsp* and *16S rRNA* genes. A different prevalence of the two *Wolbachia* sequence variants identified in *A. fraterculus* populations was observed (Table 4). We found *w*AfraCast1_A in 16% and *w*AfraCast2_A in 84% of the *A. fraterculus* individuals from our laboratory colony (37 individuals; 24 females, 13 males). In addition, we identified *w*AfraCast2_A in 95% of the insects from natural populations (39 individuals; 22 females, 17 males) while only two individuals from Puerto Yeruá (Entre Rios) showed the presence of *w*AfraCast1_A (Table 4). Based on PCR and direct sequencing, no evidence of double infections was detected in the 76 *A. fraterculus* DNA samples analyzed.

**Table 4.**
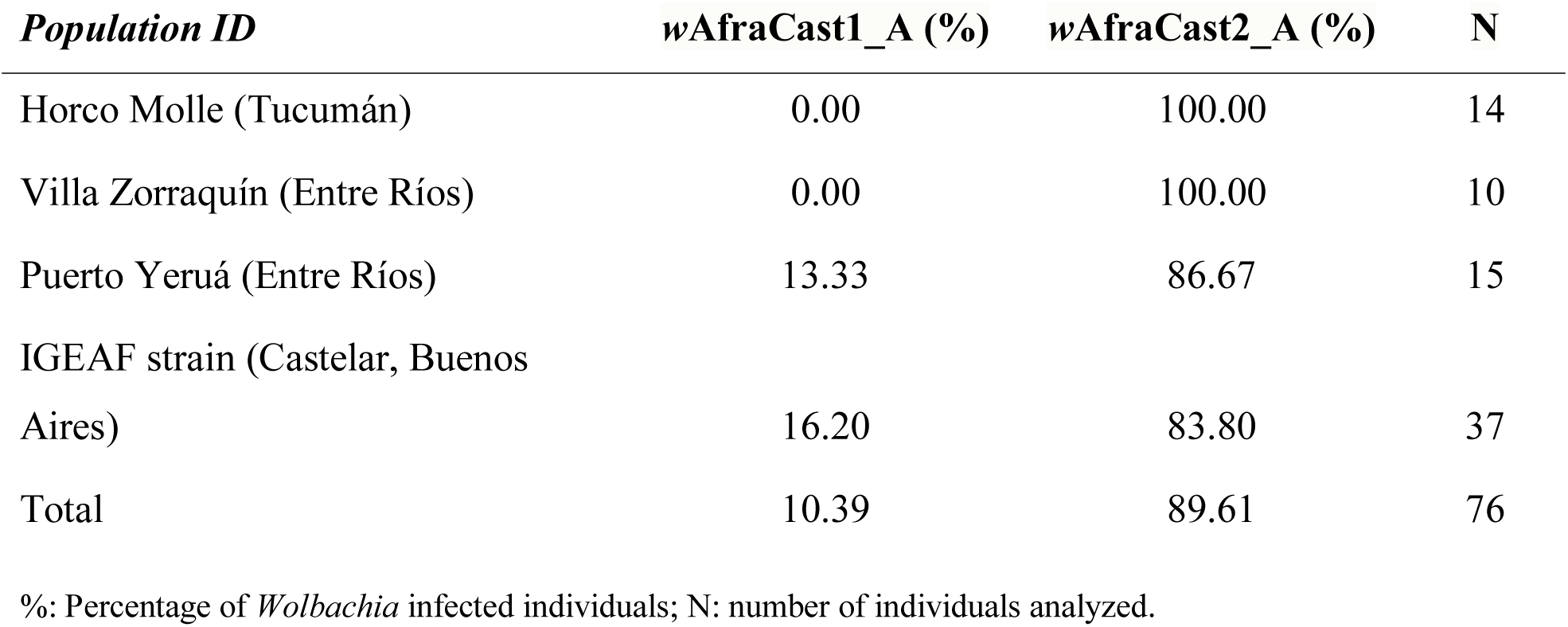
Prevalence of Wolbachia in *A. fraterculus* from Argentina.

### Cytoplasmic Wolbachia in A. fraterculus

The presence of cytoplasmic *Wolbachia* and the lack of obvious *Wolbachia* integrations into the host genome (at least detectable with the molecular methods used in the present study) were confirmed in both *A. fraterculus* strains (Af-Cast-1 and Af-Cast-2) by means of antibiotic treatment followed by PCR assays. *Wolbachia* was not detected in any of the individuals treated with antibiotic (10 flies), whereas, control individuals (10 flies belonging to Af-Cast-1 and Af-Cast-2 strains reared without antibiotic treatment) resulted in a positive *Wolbachia*-specific amplicon in 100% of the cases.

### Mating experiments

We followed the scheme of crossing experiments described in Figure 3. Parental crosses and filial crosses (sibling matings) were performed to analyze the existence of pre- and post-zygotic sexual isolation barriers associated to *Wolbachia*. Specific PCR bands of approximately 430 bp corresponding to *Wolbachia wsp* gene were successfully amplified in all *Wolbachia*-infected *A. fraterculus* individuals used in the crossing experiments (parental flies, Fig. 3). Additionally, the absence of PCR amplicons was evidenced for all *Wolbachia*-cured parental pairs used as a control of our experiments.

**Figure 3.**
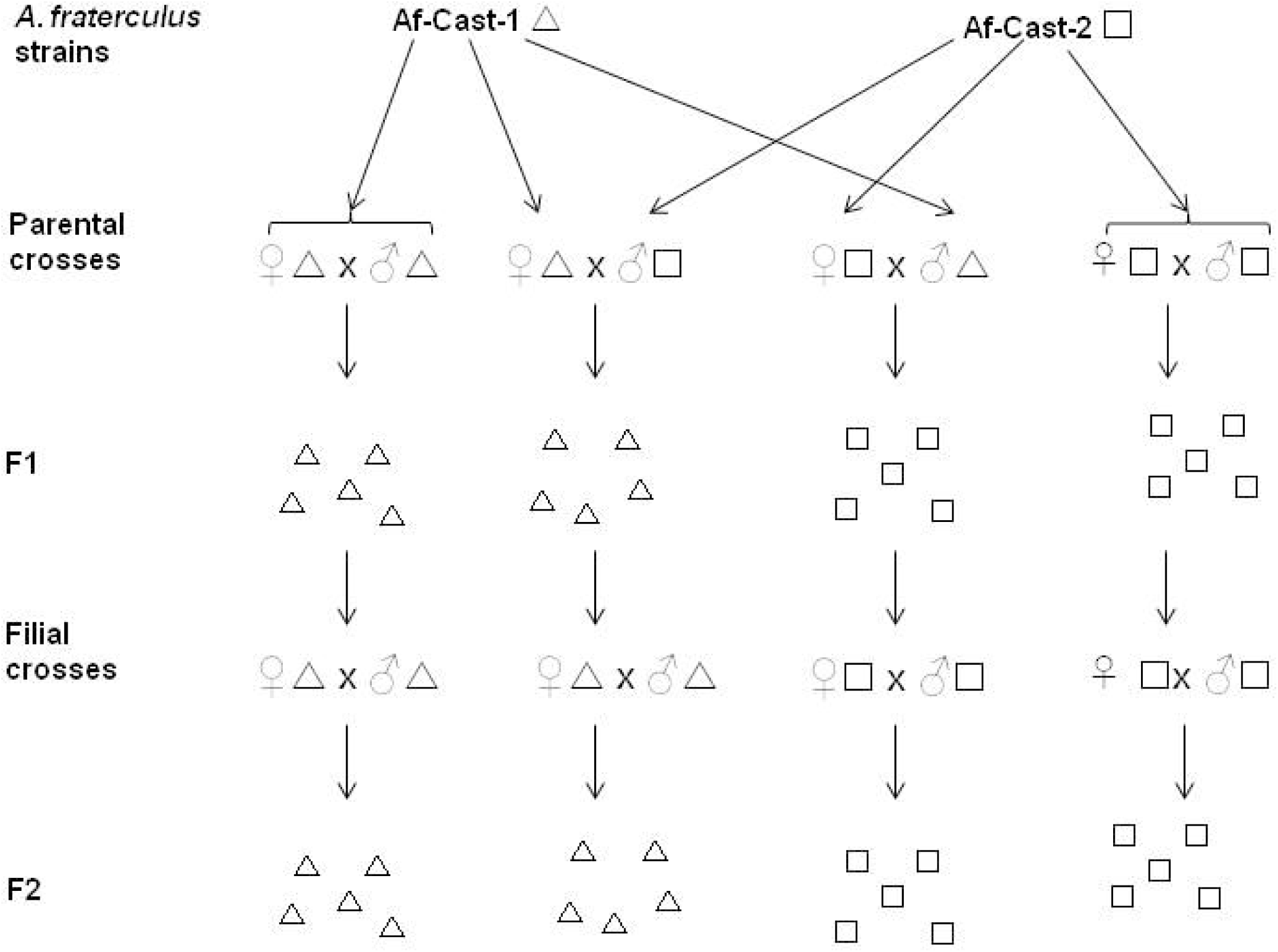
Mating scheme of Af-Cast-1 and Af-Cast-2 individuals harboriitg different variant of *Wolhachia* (wAfr aCastl_A and wAfraCast2_A, respectively). Δ *A.fraterculus* harbouring wAfraCastl_A and □ *farterculus* harbouring wAfraCast2_A. Individuals in the parental crosses were the subjects of the pre-zygotic tests. Their offspring were the subjects of the post-zygotic tests (F1).

#### Pre-zygotic isolation test

We observed similar percentages of mating among the four possible types of cross (Chi-square test: χ^2^ = 6.637, P = 0.084, d.f. = 3) with a relatively high mean percentage of mated couples (72%) compared to previous results for this species. The latency and mating duration time did not differ among the types of crosses [ANOVA: Latency: F (3,165) = 1.831, P = 0.143; Mating duration time: F (3,165) = 2.597, P = 0.054] (Table 5). These results showed a lack of pre-zygotic isolation between the *A. fraterculus sp*1 strains described here.

**Table 5.**
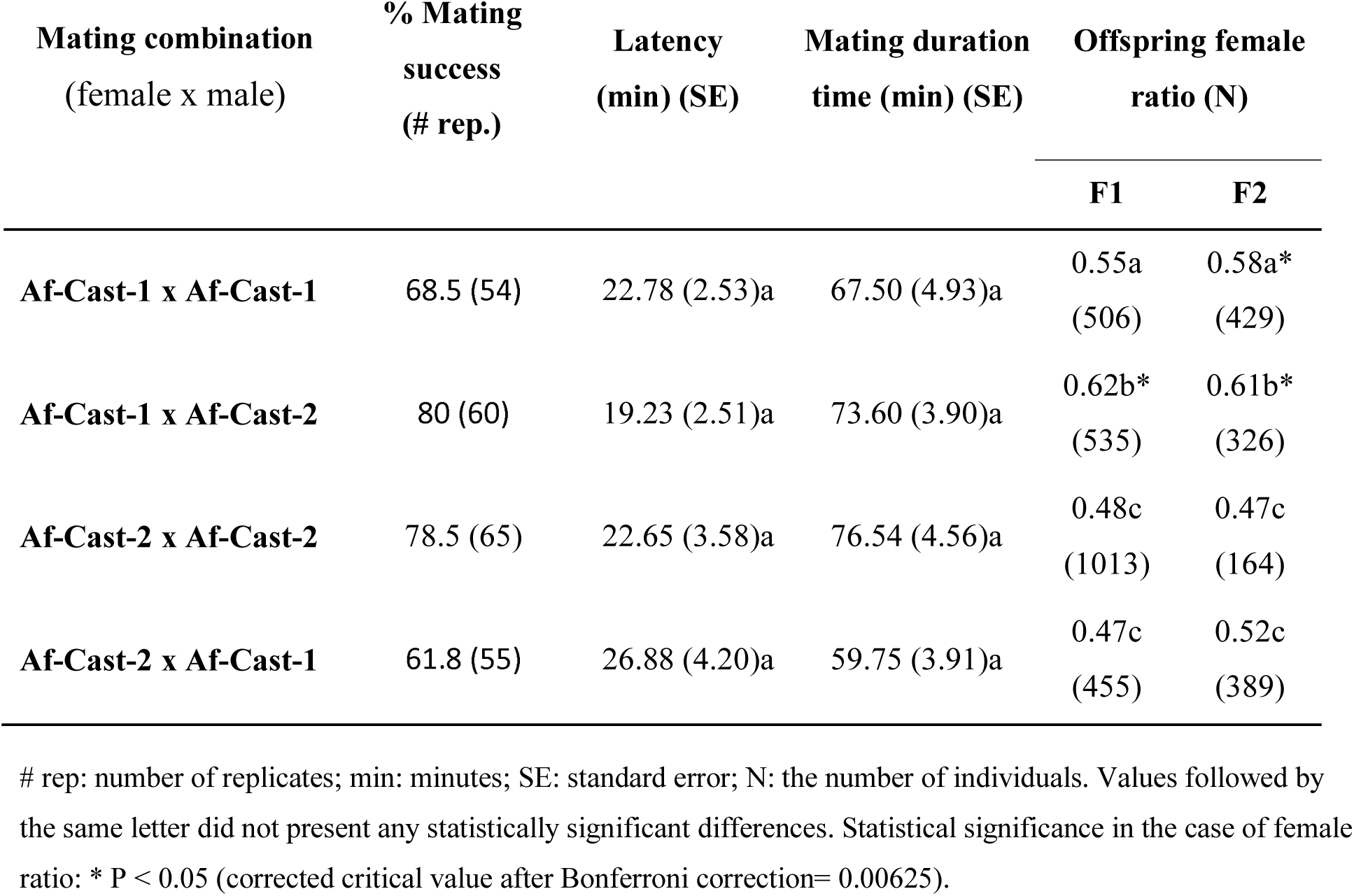
Mean values of percentage of mating, latency, and mating duration time of each type of cross, and female proportion obtained in the offspring (F1 and F2).

| Mating combination<br>(female x male) | % Mating<br>success<br>(# rep.) | Latency<br>(min) (SE) | Mating duration<br>time (min) (SE) | Offspring female<br>ratio (N) |  |
| --- | --- | --- | --- | --- | --- |
|  |  |  |  | F1 | F2 |
| **Af-Cast-1 x Af-Cast-1** | 68.5 (54) | 22.78 (2.53)a | 67.50 (4.93)a | 0.55a<br>(506) | 0.58a*<br>(429) |
| **Af-Cast-1 x Af-Cast-2** | 80 (60) | 19.23 (2.51)a | 73.60 (3.90)a | 0.62b*<br>(535) | 0.61b*<br>(326) |
| **Af-Cast-2 x Af-Cast-2** | 78.5 (65) | 22.65 (3.58)a | 76.54 (4.56)a | 0.48c<br>(1013) | 0.47c<br>(164) |
| **Af-Cast-2 x Af-Cast-1** | 61.8 (55) | 26.88 (4.20)a | 59.75 (3.91)a | 0.47c<br>(455) | 0.52c<br>(389) |

#### Post-zygotic isolation analysis

We did not observe any statistically significant differences among the types of crosses regarding the percentage of eggs that hatched and adults that emerged in the F1 generation [%Egg hatch: F (3,82) = 0.52, P = 0.67; % Adults emergence: F (3,48) = 0.28, P = 0.84]. In contrast, the percentage of pupation showed statistically significant differences among crosses [ANOVA: F (3,46) = 4.78, P < 0.01]. Multiple comparison analysis showed that the Af-Cast-1 x Af-Cast-1 cross had a statistically significant lower percentage of pupation than the Af-Cast-2 x Af-Cast-2 cross. The other two types of crosses (Af-Cast-1 x Af-Cast-2 and Af-Cast-2 x Af-Cast-1) showed intermediate pupation values (Fig. 4 A-C).

**Figure 4.**
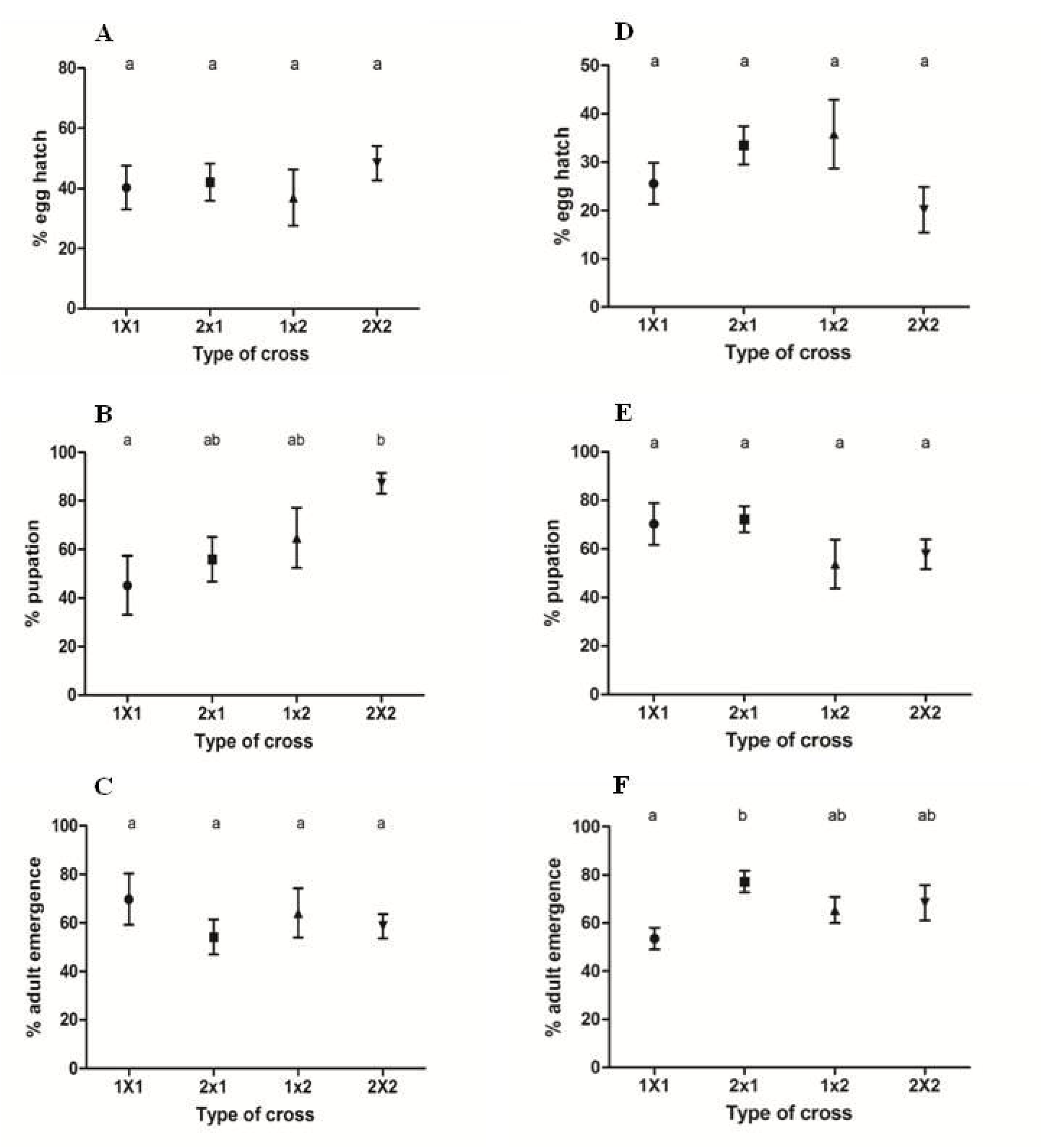
Survival across development. Parameters measured for each type of cross (female x male). The crosses Af-Cast-1 x Af-Cast-1, Af-Cast-2 x Af-Cast-1, Af-Cast-l x Af-Cast-2, Af-Cast-2 x Af-Cast-2 are mentioned in the figure as l x 1, 2 x 1, l x 2 and 2 x 2 respectively. A, B and C - Fl offspring analysis. D, E and F-F2 offspring analysis. (A/D) mean (± S.E.) % egg hatch; (B/E) mean(± S.E.) %pupation = percentage of larvae that reached pupae stage; (C/F) mean(± S.E.) % adult emergence = percentage of pupae that reached the adult stage. Points sharing a letter did not present any statistically significant differences.

In the F2 generation, we observed that the percentage of egg hatch and the percentage of pupation showed no statistically significant differences among crosses [F (3,30) = 2.15, p = 0.18; and F (3,29) = 1.49, p = 0.24, respectively] (Fig. 4 D and E). However, the percentage of adult emergence showed statistically significant differences among crosses (F (3,28) = 3.46; p = 0.029). Furthermore, Af-Cast-1 x Af-Cast-1 families showed the lowest percentages of adult emergence and Af-Cast-2 x Af-Cast-1 families the highest (Tukey test) (Fig. 4 F).

A sex ratio distortion that significantly favored females (both in F1 and F2 offspring) was detected in Af-Cast-1 x Af-Cast-2 crosses, whereas, in the case of Af-Cast-1 x Af-Cast-1 crosses, significant deviation of this parameter was observed only in F2 descendants. No bias was evidenced in crosses involving Af-Cast-2 females (Table 5).

Further analysis of data obtained from the parental crosses gave no statistically significant differences regarding percentage of mated females that produce eggs (χ^2^ = 2.321; p=0.508, d.f. = 3), percentage of females that produce viable eggs (χ^2^ = 2.322, p = 0.508, d.f. = 3), percentage of females with descendants (χ^2^ = 0.396, p = 0.941, d.f. = 3), percentage of females that produce viable eggs (χ^2^ = 4.893, p = 0.180, d.f. = 3) and percentage of females with descendants (χ^2^ = 5.778, p = 0.123, d.f. = 3), (Fig. 5 A-E). Since data were homogeneous, results were pooled and compared between types of female. Again, the percentage of mated females that produced eggs did not differ between type of female (χ^2^ = 1.956, p = 0.162, d.f. = 1) (Fig. 5 F). Similarly, the percentage of females that produce viable eggs and the percentage of females with descendants were not statistically different between types of female (χ^2^ = 0.632, p = 0.427, d.f. = 1 and χ^2^ = 0.070, p = 0.791, d.f. = 1, respectively) (Fig. 5 G and H). In contrast, both the percentage of mated females that produced viable eggs and the percentage of mated females with descendants were significantly higher for the Af-Cast-2 females (χ^2^ = 4.706, p = 0.030, d.f. = 1; and χ^2^ = 5. 560, p = 0.018, d.f. = 1, respectively) (Fig. 5 I and J).

**Figure 5.**
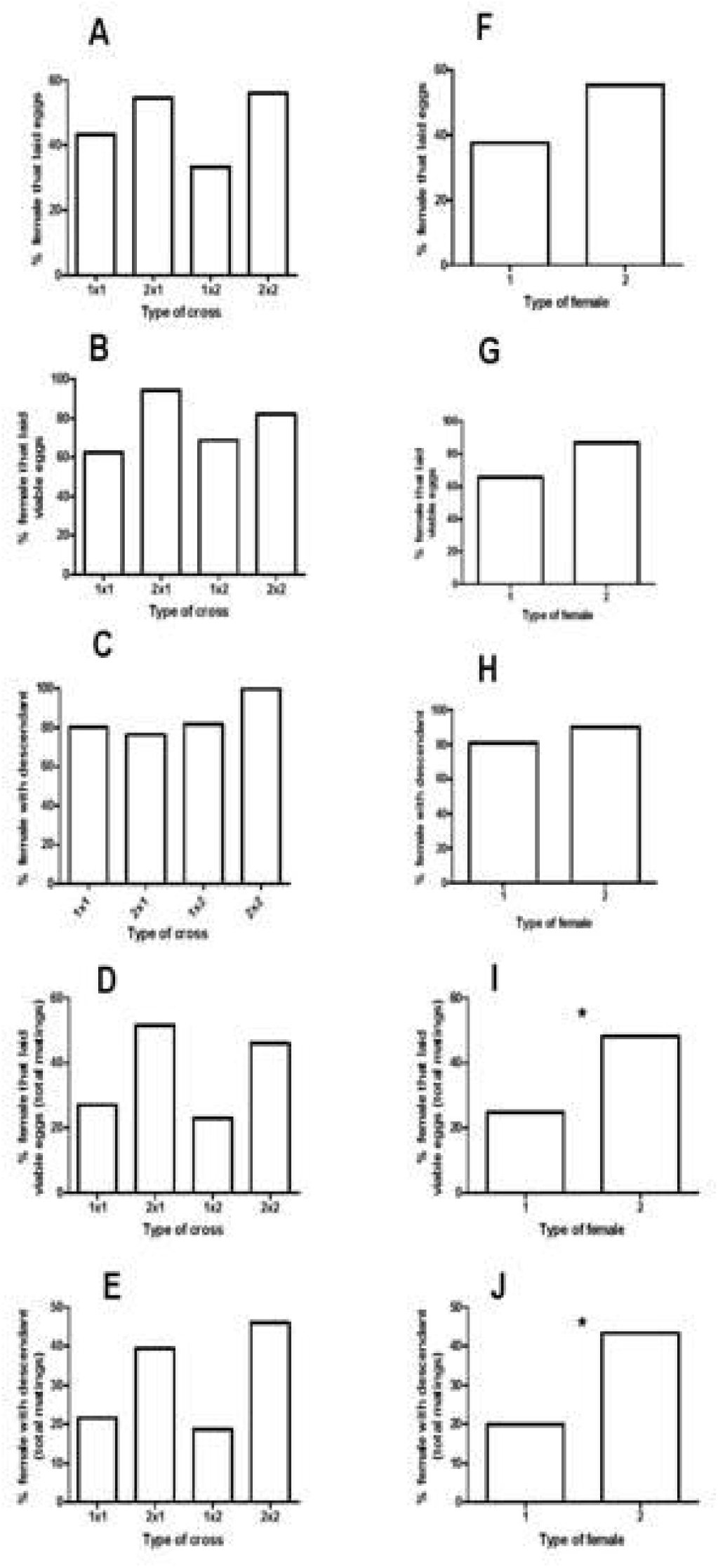
Mating experiments - additional analyses. **A-E** represent comparisons that included the four types of crosses. **F-J,** datacoming from the same female were pooled irrespectively of the type of male they mated. Asterisks indicate statistically significant differences (p < 0.05) when percentages were compared by means of a Chi-square test of homogeneity.

### Cytology of mated females

For each type of cross, we dissected the ovaries of at least 10 mated females that did not lay eggs and five mated females that laid unviable eggs. In all cases (77 females), we observed ovaries with a normal shape (fully developed and conserved size and structure), similar to the ones observed in reproductively mature females (control females, 15-20 days old) from *A. fraterculus* IGEAF strain (data not shown). In addition, the cytological analysis of spermathecae showed a high density of sperm (bundles) present in control females (Fig. 6 A) and absence of sperm in females that did not lay eggs and females that produced unviable eggs from the crossing experiments (77 females analyzed) (Fig. 6 B). It is worth mentioning that *A. fraterculus* is able to lay unfertilized eggs even in the absence of mating (virgin females). The results obtained here highlight the absence of sperm in the spermathecae as the main cause of the lack of descendants in the analyzed crosses.

**Figure 6.**
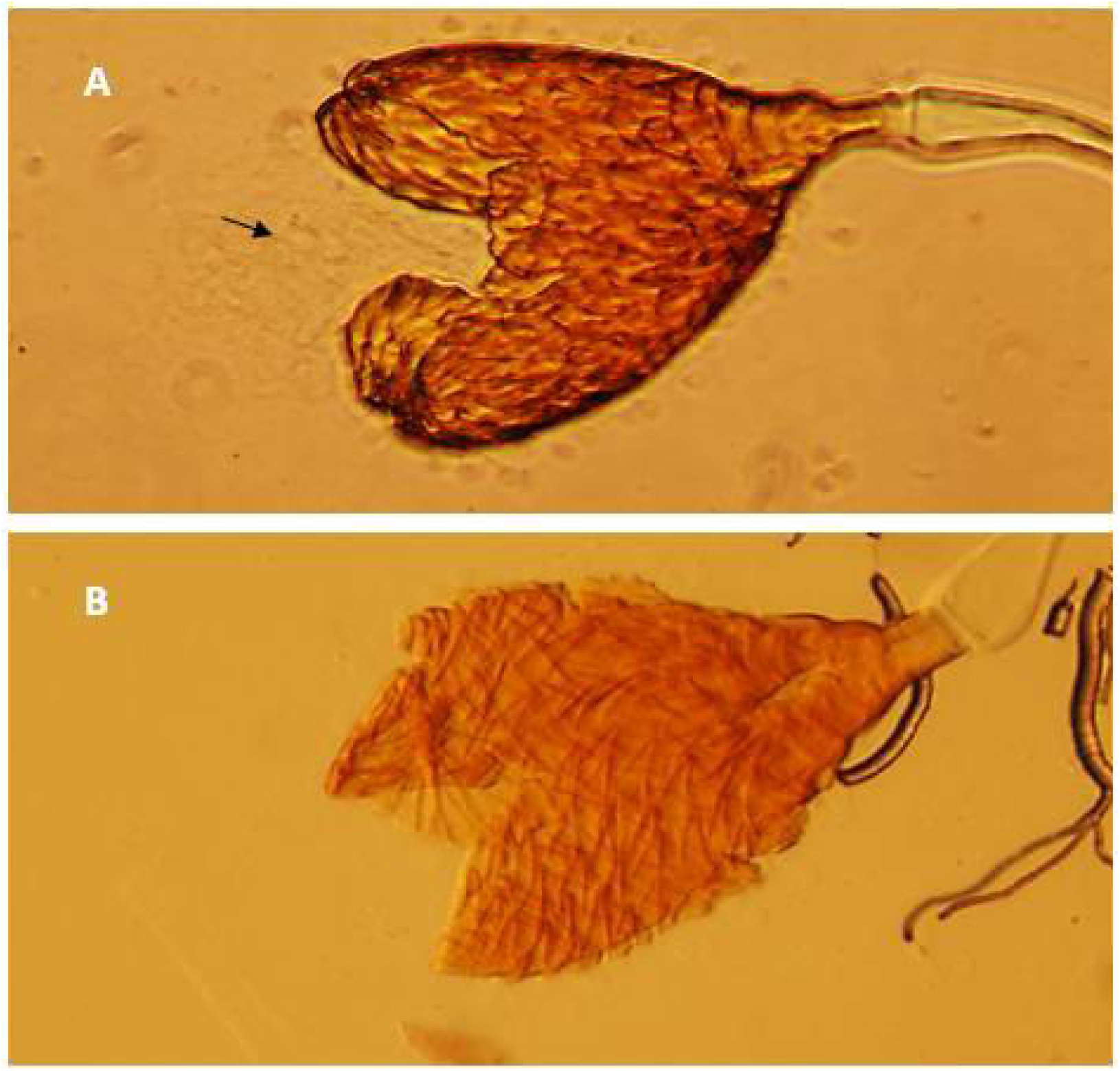
Cytological an alysis of A.fraterculus spen natb ecae (20X). A: spermatheca of *A.fraterculus* showing the presence of sperm bundles, which are indicated by an arrow **B:** spermatheca of *A.fraterculus* showing no sperm in its content.

### Detection of other reproductive symbionts

We evaluated the presence of *Spiroplasma sp*., *Cardinium sp., Rickettsia sp.*, *Arsenophonus sp.* and *Hamiltonella sp.* by using specific PCR assays (Table 2). After the analysis of at least ten DNA samples from each of the *A. fraterculus* IGEAF strains, no symbiont-specific amplicons were obtained.

## DISCUSSION

The presence of *Wolbachia* in both laboratory and wild *A. fraterculus* populations from Argentina was evidenced and characterized in this study. Mating experiments showed a slight deficit of males in F1 and F2 progenies and a detrimental effect on larval survival, suggesting that some kind of male-killing phenotype may be associated with the presence of one of the two *Wolbachia* strains detected in *A. fraterculus sp.* 1.

The analysis of the *wsp* gene at a nucleotide level allowed the identification of two sequence variants of *Wolbachia* in the host populations (named as *w*AfraCast1_A and *w*AfraCast2_A). Sequence analysis of concatenated MLST dataset showed that these *Wolbachia* variants share the same MLST allelic profile. Furthermore, phylogenetic analysis clustered these variants in the same group (ST13) with *w*Mel (*Wolbachia* infecting *D. melanogaster*), along with other *Wolbachia* strains belonging to supergroup A. Our findings using MLST in the identification of *Wolbachia* (and its clustering in supergroup A) were also supported by *16S rRNA* sequence analysis.

Further characterization of *Wolbachia* using an antibiotic treatment, allowed the confirmation of an active cytoplasmic infection of this endosymbiont. We did not find evidence of insertion in the *A. fraterculus* genome, as antibiotic-treated flies showed lack of a specific amplicons for *wsp* and *16S rRNA Wolbachia* genes. In addition, the prevalence analysis of the *Wolbachia* variants shows the absence of double infections under the experimental design and standard conditions used in the present study. Single infections of *Wolbachia* have also been described in other *A. fraterculus* populations [79, 117, 118].

The presence of *Wolbachia* in Argentinean populations of *A. fraterculus* was first reported by Cáceres et al. [79]. These authors analyzed two laboratory strains of *A. fraterculus* established at Insect Pest Control Laboratory (Seibersdorf, Austria), originally derived from wild flies collected from Argentina and Peru. Each laboratory population harbored closely related *Wolbachia* strain (*w*Arg and *w*Per, respectively), with the presence of one nucleotide substitution in *w*Arg based on *wsp* gene sequencing. In the present work, we found identical results at a nucleotide level with these previously reported *Wolbachia wsp* gene sequences (*w*AfraCast1_A identical to *w*Per and *w*AfraCast2_A identical to *w*Arg). Moreover, we found that the *w*AfraCast1_A *wsp* sequence presented an identical nucleotide composition compared to a partial *wsp* sequence detected in a Brazilian population of *Anastrepha sp*.1, (GenBank AN EU 116325) reported by Coscrato and colleagues [117]. The presence of the same *wsp* gene sequence in different populations of *A. fraterculus* does not necessarily mean that they are infected with identical *Wolbachia* strains [16, 119]. The *Wolbachia* infection status of several morphotypes of the *A. fraterculus* cryptic species complex (including *A. fraterculus sp.*1) was recently published by Prezotto et al. [93]. The information provided by these authors with respect to *Wolbachia* sequence variants infecting different *A. fraterculus* populations from Argentina (either using MLST or *wsp* HVR analyses) differs from our findings. More knowledge regarding the origin of the samples and the number of individuals analyzed by Prezotto et al. [93] are needed in order to compare the results obtained in the two studies. Moreover, the same authors suggested a potential association between specific *Wolbachia* strains and distinct *A. fraterculus* morphotypes, which could act as a reinforcing factor in the diversification processes, providing also, some evidence of the possible way of transmission of *Wolbachia*. Further characterization of *Wolbachia* strains infecting members of the *A. fraterculus* complex, taking into account crossing experiments and deeper molecular analysis could provide insight to the speciation process in this complex, unraveling the genetic entities present and their phylo-geographic distribution.

Our crossing experiments showed a detrimental effect during the development for crosses involving Af-Cast-1 females. This is suggested by a statistically significant lower percentage of pupation in F1 offspring and a lower percentage of adults emergence in F2 descendants observed in the crosses involving Af-Cast-1 flies. Despite the lack of differences between females in the percentage of mated females that laid eggs, those that laid viable eggs, and those that successfully produced progeny, we were able to find a tendency to lower values in Af-Cast-1 females, which was statistically significant when these percentages were computed considering the total number of mated females, which allowed these small, non-significant effects, to accumulate. These results might point to a negative effect of a *Wolbachia* variant on the reproductive biology of its host. We also found that some parameters associated to immature development varied in some crosses between F1 and F2. For instance, Af-Cast-2 x Af-Cast-2 cross yielded higher egg hatch and pupation in the F1 than in the F2. Because these crosses involved flies with equivalent genetic background and *Wolbachia* infection status, this result suggests that unidentified experimental conditions probably varied between the F1 and F2.

Cytological analysis showed the absence of sperm in the spermathecae of females that did not lay eggs and females that produced unviable eggs, showing that the lack of sperm transfer is the main cause of unviable embryos production in some families. This result combined with the lack of differences in the % of hatched eggs allowed us to rule out the presence of a bidirectional cytoplasmic incompatibility associated to *Wolbachia* infection in the tested crosses. Also, it supports the hypothesis that detrimental effects in the survival associated to *Wolbachia* would occur later in the developmental stages rising new questions regarding possible effects of this bacterium on the host’s reproductive behavior that should be further addressed.

The analysis of sex ratio in each type of cross and generation showed a distortion in favor of females in crosses involving Af-Cast-1 females. Particularly, we observed this type of distortion in F1 and F2 of Af-Cast-1 x Af-Cast-2 pairs, and F2 offspring from Af-Cast-1 x Af-Cast-1 crosses. Additionally, individual analyses of each family showed that only a few paired-crosses appear to contribute with this sex ratio distortion (Additional File 3).

Our finding indicates that the effect of *Wolbachia* may not be homogeneous among different individuals belonging to the same host strain and requires further analysis. Studies including the quantification of *Wolbachia* titers in the parental couples and the measurement of biologically important parameters, in connection with genetic studies of the offspring, including cytological (cytogenetic) analysis will provide more evidence of the phenotype elicited by this endosymbiont in *A. fraterculus.* In this regard, previous studies described the importance of bacterial densities in the expression of a phenotype and the presence of different *Wolbachia* densities during host development [2, 36] using sensitive tools as the quantitative real time PCR (qPCR) and other methods for the detection of low titer reproductive symbionts [120–124]. Moreover, the action of non-bacterial, maternal-inherited microorganisms [125] must also be taken into account for future studies. Detection of low titer endosymbionts using more sensitive methods and the inclusion of crossing experiments involving antibiotic treatments will contribute to a better understanding of our findings.

Detrimental effects (lower % of pupation and % of adult emergence in the F1 and F2, respectively) and the sex ratio distortion observed in crosses involving Af-Cast-1 females, potentially elicited by the presence of *Wolbachia* and associated to a male killing phenotype, have been previously described in insect species by Hurst et al. [45], Dyer and Jaenike [46] and Kageyama and Traut [126]. A larger set of crossing experiments combined with the analysis of several biologically important parameters from the host populations (e.g. fecundity, % egg hatch, and/or differences in larval and/or pupal survival) are needed to better understand the effects *Wolbachia* may be inducing to this host species.

The results obtained here display the differences between the phenotype elicited by two *Wolbachia* sequence variants on their hosts, revealing some disparity in the cross-talk involving the bacteria and its hosts. This may include genetic variability in the bacterium as well as in the host species. In our study, we evidenced a significant similarity between the two *Wolbachia* strains analyzed, based on the identical MLST allelic profile and the identical sequences of *16S rRNA* gene and seven additional *Wolbachia* genes (*gro*EL, *glt*A, *dna*A, *suc*B, *asp*C, *atp*D and *pdh*B). It is also worth noting that several studies have demonstrated the importance of the host genetic background associated to the molecular mechanisms involved in the phenotype induced by *Wolbachia* [39, 58, 118, 127]. Microsatellite analyses have shown a high genetic variability and differentiation among Argentinean populations of *A. fraterculus* [90, 128, 129]. Genetic evaluations using this kind of markers could be potentially useful to identify variation between the *A. fraterculus* strains harboring different *Wolbachia* variants under study in the present work. These studies may contribute to our understanding of the different reproductive effects displayed by *Wolbachia* in these singly-infected *A. fraterculus* strains.

### CONCLUSION

This work contributes to the characterization of *Wolbachia* infection in *A. fraterculus sp*.1 from Argentina. We gained a first insight on possible mechanisms associated to the *Wolbachia* - *A. fraterculus* interaction by crossing singly-infected *A. fraterculus* strains. We found a potential deleterious effect on immature stages and a sex ratio distortion (male-killing) associated to one of the detected *Wolbachia* variants (*w*AfraCast1_A). Further mating experiments, coupled with quantification of *Wolbachia* titers and including cured lines, will shed light on the phenotype elicited by *Wolbachia* in *A. fraterculus*. Our findings are important for the characterization of *A. fraterculus* populations from Argentina, and as a contribution to develop environmentally-friendly and species-specific control strategies against this pest.

## Ethics approval

Not applicable.

## Consent for publication

Not applicable.

## Availability of data and materials

*Wsp* gene sequences generated in this study from *w*AfraCast1_A and *w*AfraCast2_A have been deposited in the *Wolbachia* MLST (pubmlst.org/*Wolbachia*/) and GenBank, National Center for Biotechnology Information. (NCBI) databases under accession numbers KC589026.1 and KC589027.1. Allelic profile of MLST scheme of five genes (*gat*B, *cox*A, *hcp*A, *fbp*A and *fts*Z) from *w*AfraCast1/2_A and, HVRs allelic profile of *w*AfraCast1_A and *w*AfraCast2_A are available on *Wolbachia* MLST database.

Nucleotide sequences of the *gro*EL, *glt*A*, dna*A*, suc*B*, asp*C*, atp*D and *pdh*B genes from *Wolbachia* infecting Argentinean *A. fraterculus* were submitted to GenBank (https://www.ncbi.nlm.nih.gov/genbank/index.html) under accession numbers MG977022-28 respectively.

Raw data obtained in this work is available upon request to the corresponding author.

## Competing interests

The authors declare that they have no competing interests.

## Funding

This work was supported by funds from the National Institute of Agricultural Technology (INTA) through the projects PNBIO 11031023 and AEBIO-242411 (module pests) to SBL; the International Atomic Energy Agency through the Research Contract N°17041 to DFS and the Agencia Nacional de Promoción Científica y Tecnológica (Argentina) through the project Foncyt-PICT 2013-0054 to DFS. CAC was supported by the Post-grade and Retraining Program from INTA (2014).

## Authors’ contributions

CAC and FHM maintained the insects and conducted experimental assays. SBL, DFS, KB and JLC conceived the study. CAC and AAA performed symbiont’s identification. CAC performed all molecular assays and analysis. CAC, SBL, FHM and DFS conducted mating assays. SBL, DFS, KB and JLC conducted the data analysis, and wrote the manuscript. All authors read and approved the final manuscript.

## Acknowledgements

Authors are grateful to María Teresa Vera (University of Tucumán, Argentina) and Juan Pedro Bouvet (INTA Concordia) for their invaluable help in the sampling of infested fruits. Also authors are indebted to Carlos Cáceres and the IPLC Seibersdorf staff (FAO/IAEA, Vienna, Austria) for their assistance in an important part of the activities of this project. We thank the anonymous reviewers for their constructive comments, which helped us to improve the manuscript.

